# Cas9AEY (Cas9-facilitated Homologous Recombination Assembly of non-specific Escherichia coli yeast vector) method of constructing large-sized DNA

**DOI:** 10.1101/2024.09.17.611575

**Authors:** xiaoshu ma, lei yang, hua ye

## Abstract

Saccharomyces cerevisiae is widely used in DNA assembly due to their efficient homologous recombination [1], but DNA assembly through yeast recombination in vivo usually requires the vector to have the ability to replicate in yeast. The CRISPR-Cas9 system can efficiently edit DNA [2,3], and the system can also be used for DNA editing of plasmids. In this paper, a yeast universal element is selected, which can be inserted into the vector, so that the vector can replicate in yeast cells, and then the intermediate plasmid containing yeast universal element can be obtained by recombination in yeast. At the same time, a pCas-SmR plasmid was designed in this paper. After Donor DNA is added, the CRISPR-Cas9 system can accurately and efficiently knock out the yeast universal element in the intermediate plasmid, remove the pCas-SmR plasmid through sucrose screening, and finally obtain a pure plasmid. Saccharomyces cerevisiae cells are widely used in DNA assembly due to their efficient homologous recombination [1], but DNA assembly through yeast recombination in vivo usually requires the vector to have the ability to replicate in yeast. The CRISPR-Cas9 system can efficiently edit DNA [2,3], and the system can also be used for DNA editing of plasmids. In this paper, a yeast universal element is selected, which can be inserted into the vector, so that the vector has the ability to replicate in yeast cells, and then the intermediate plasmid containing yeast universal element can be obtained by recombination in yeast. At the same time, a pCas-SmR plasmid was designed in this paper. After Donor DNA is added, the CRISPR-Cas9 system can accurately and efficiently knock out the yeast universal element in the intermediate plasmid, remove the pCas-SmR plasmid through sucrose screening, and finally obtain a pure knocked out plasmid.

## Introduction

With the rapid development of the field of gene synthesis, especially the in-depth research in the field of genome in recent years, researchers have increasingly high requirements for the assembly technology of large fragments of DNA [4]. The usual de novo synthesis process of genes includes chemical synthesis of oligonucleotides below 100nt, assembly of oligonucleotides into double-stranded DNA of 500bp to 1000bp based on polymerase chain reaction [5], and finally splice into final DNA by long fragment DNA assembly technology [6]. At present, the methods used for synthetic synthesis of large fragments of DNA mainly include in vitro assembly and in vivo assembly. Commonly used in vitro assembly methods include traditional enzyme ligand technology, Gibson assembly technology based on exonuclease, Golden Gate technology based on endonuclease, etc. [7,8]. Among them, the isothermal one-step splicing technique [9] invented by Gibson is more commonly used, which can splice multiple DNA fragments with overlapping regions together in one step. The host cells selected by the in vivo assembly method mainly include Escherichia coli, Saccharomyces cerevisiae, Bacillus subtilis and other model organisms, among which yeast in vivo homologous recombination is commonly used. Saccharomyces cerevisiae is used for the synthesis of large fragments of DNA due to its efficient homologous recombination ability. At present, researchers have used yeast in vivo homologous recombination to complete megatons of genome synthesis [10].

In order to replicate in yeast cells, the vector used for yeast splicing must contain yeasty-related promoters, such as yeast centromere elements, autonomous replication sequences CEN/ARS, 2u promoter, etc. [11,12], and most plasmids in yeast cells are single copies, which is difficult to extract, and it is usually necessary to shuttle to Escherichia coli for plasmid enrichment. Therefore, the vector also needs to contain promoters that can replicate in E. coli, so the use of yeast splicing technology is generally limited to the vector.

CRISPR/Cas9 technology is a breakthrough in the biological field in recent years, providing a more efficient method for gene editing [13]. CRISPR is a repetitive DNA sequence. When bacteria are invaded by viruses, they can store part of the DNA of viruses into the CRISPR region. When viruses invade again, bacteria will cut the DNA of viruses according to the stored sequence [14]. Cas gene is CRISPR-related gene, which can work together with CRISPR to break the target DNA double strand [15]. Using CRISPR/Cas9 technology, genes can be knocked in and knocked out.

Traditional yeast splicing to assemble large fragments of DNA requires the use of shuttle vectors, although the success rate is high, but its application is severely limited. In this paper, we present a simple and rapid method for the synthesis of plasmids of 30kb and above. The method is mainly based on yeast in vivo recombination technology and CRISPR/Cas9 technology. Firstly, a gene sequence, called yeast universal element, was designed to enable the recombinant plasmid to replicate and pass through saccharomyces cerevisiae cells, and the intermediate plasmid containing yeast universal element was obtained by using the sequence for yeast homologous recombination. Finally, the final plasmid was obtained by knocking out the yeast universal element with CRISPR/Cas9 technology. Here we describe in detail the design of yeast universal element and tool plasmid pCas-SmR. The yeast universal element designed in this study was 1.8kb in length and mainly consisted of two functional regions. The optimized CEN/ARS region could ensure the replication of the recombinant plasmid in yeast cells [16,17], and the URA region could enable the yeast strain to grow in the medium lacking uracil [18]. The tool plasmid pCas-SmR mainly consists of Cas functional region that plays a role in DNA cutting, λRed homologous recombination system induced by arabinose [19], sgRNA sequence of yeast universal element and SacB sucrose screening system [20]. The role of this tool plasmid is to eliminate yeast universal element accurately and efficiently. Firstly, fragments of DNA, linearized vectors and yeast universal elements were transformed into Saccharomycetales cells by lithium acetate [21], and positive clones were screened by uracil defect medium to obtain intermediate plasmids containing yeast universal elements. By electrical transformation, the intermediate plasmid and the designed donor DNA fragment are co-transformed into Escherichia coli expressing pCas-SmR plasmid, which can complete the knockout of the yeast universal element and obtain the final construction. Finally, pCas-SmR plasmid was deleted by sucrose screening.

## Result

### Recombinant splicing intermediate plasmids in yeast

In order to verify the splicing efficiency of yeast universal elements, we tested the assembly experiments of 14kb and 32kb plasmid. Among them, the vector of 14KB plasmid was PCDNA3.1, and the vector of 32KB plasmid was adenovirus vector. The 14kb plasmid was divided into two 4.5kb insertion fragments and a 5kb carrier fragment, and the 32kb plasmid was divided into two 10.5kb insertion fragments and 11kb carrier fragments. Each fragment was synthesized by conventional methods in Suzhou genwiz Company, and each fragment and yeast universal element were converted into Saccharomyces cerevisiae BY4741 by lithium acetate. primers were designed to perform colony PCR detection at fragment junctions. The PCR fragment sizes at the three junctions of the 14KB plasmid were 2330bp at the No. 1 junction, 980bp at the No. 2 junction, 668bp at the No. 3 junction, and the PCR fragment sizes at the three junctions of 32KB plasmid were: Junction 1 is 2037bp, junction 2 is 1637bp and junction 3 is 401bp. According to the PCR detection results of yeast colonies (Fig1), all fragment junctions could be detected by PCR, and the positive rate of yeast splicing of 14kb plasmid and 32kb plasmid was 100%. That is, the yeast universal element designed in this study can efficiently assemble large fragments of non-specific E. coli vectors in saccharomyces cerevisiae cell BY4741.

**Fig1:**
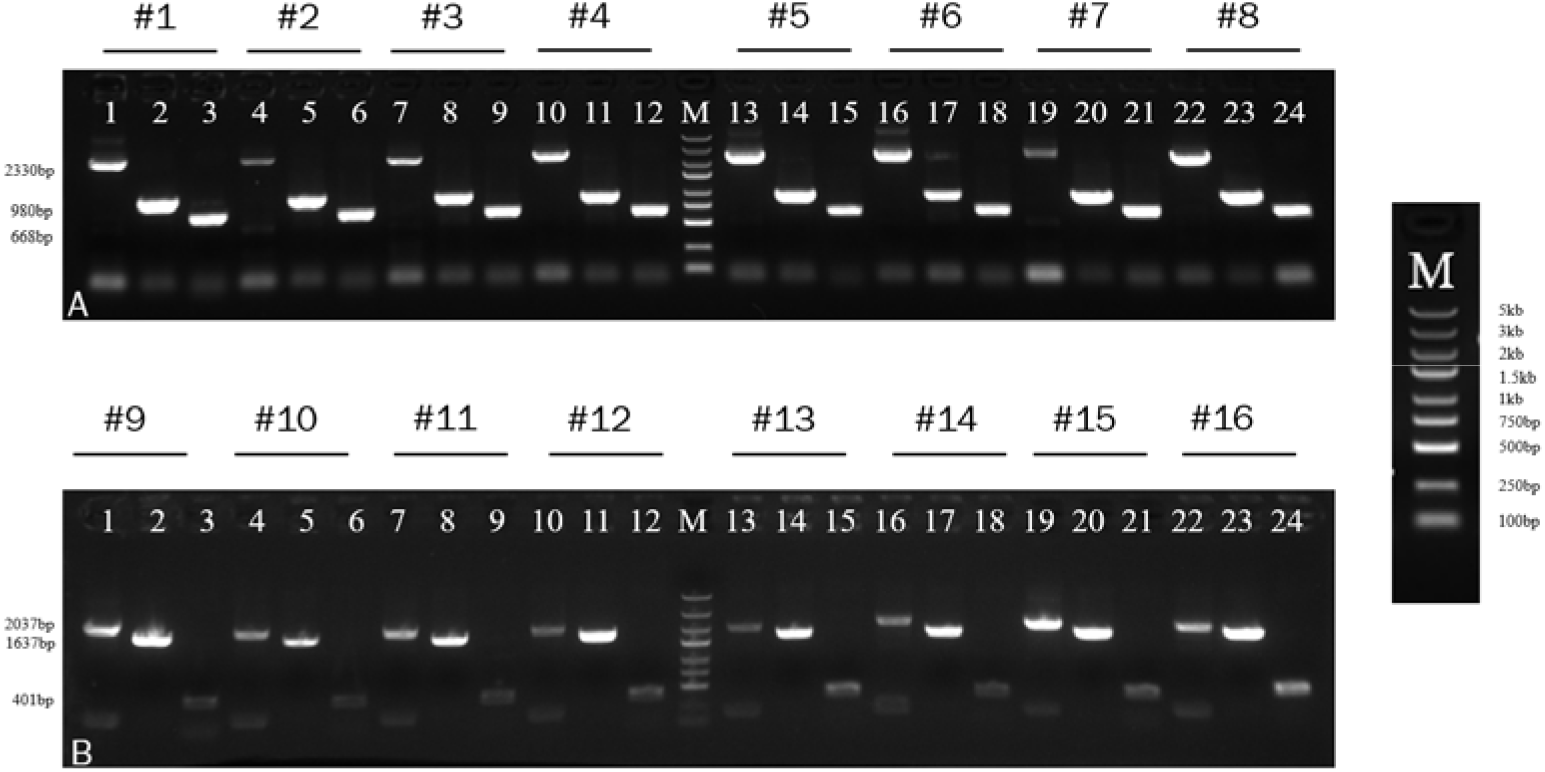
Single colony PCR detection of yeast colonies 8 single colonies were selected from each of the 14kb (#1∼#8) and 32kb (#9∼#16) plasmids for colony PCR detection. (A) The PCR results of 14kb plasmid colonies showed that lane 1-3 was the No. 1-3 connector of #1 colony, and so on, and lane 4-6 was corresponding to the No. 1-3 connector of #2 colony, so it is not necessary to go into details one by one. M is 5000 bp DNA Marker. (B) The PCR results of 32kb plasmid colonies showed that lane 1-3 was the No. 1-3 connector of #9 colony, and so on, and lane 4-6 was corresponding to the No. 1-3 connector of #10 colony, and so on

### Yeast universal element knocked out

To eliminate the influence of yeast universal element on the subsequent application of the plasmid, we used CRISPR/Cas9 technology to knock out yeast universal element. First, we constructed a tool plasmid pCas-SmR specially used to knock out yeast universal element, and the plasmid map is shown in Figure 2. At the same time, a 900bp Donor DNA was amplified by fusion PCR using the 450bp sequence adjacent to the yeast universal element as the template. The plasmid to be knocked and Donor DNA were co-transformed into Escherichia coli expressing pCas-SmR plasmid to knock out the yeast universal element. PCR identification of the colony after knockout showed that the size of the successful knockout of 14KB plasmid was 1663 bp, the size of the failed knockout was 3495 bp, the size of the successful knockout of 32KB plasmid was 1306bp, and the size of the failed knockout was 3138bp. As can be seen from the figure (Fig3A, D) All PCR bands of the selected clones were successfully knocked out, which proved that the yeast universal element regions of the selected clones were successfully knocked out. The plasmid was verified by enzyme digestion, as shown in Figure 3 B, C. (Figure 3 B, C) showed the results of enzyme digestion of 14 kb plasmid. The strip size of the original plasmid was 9387 bp, 4848 bp and 1838 bp. After the deletion of yeast universal element, the plasmid size was 9387 bp, 4848 bp, 6 bp. (Fig3E, F) was the verification result of the enzyme digestion of 32kb plasmid, and the size of the enzyme digestion result of the original plasmid was 10973 bp, 9207 bp, 6715 bp, 5520 bp, 1838 bp. The results of plasmid digestion were 10973 bp, 9207 bp, 6715 bp, 5520 bp, 6 bp after knocking out the yeast universal element. Compared with the results before knockout, there was no band of yeast universal element in the results after knockout. The plasmid with the yeast universal element knocked out was subjected to ONT third-generation sequencing. Using the plasmid sequence with the yeast universal element not knocked out as the template, it was obvious that the yeast universal element was missing, and the reading value of this region tended to 0 compared with Coverage of other regions, while the rest of the sequences were not affected. The results showed that the yeast element was successfully knocked out.

**Fig2.**
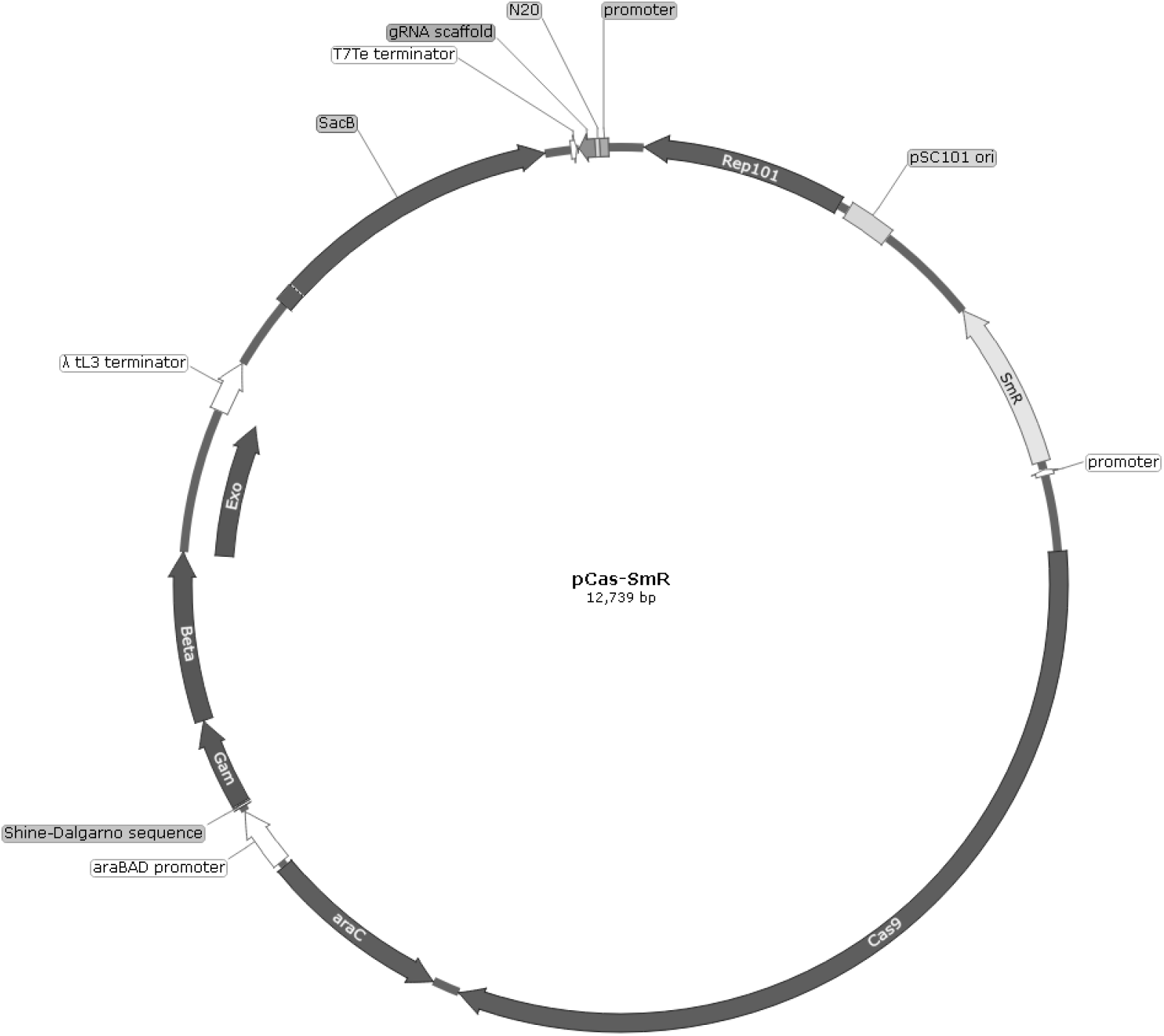
pCas-SmR plasmid map

**Fig3.**
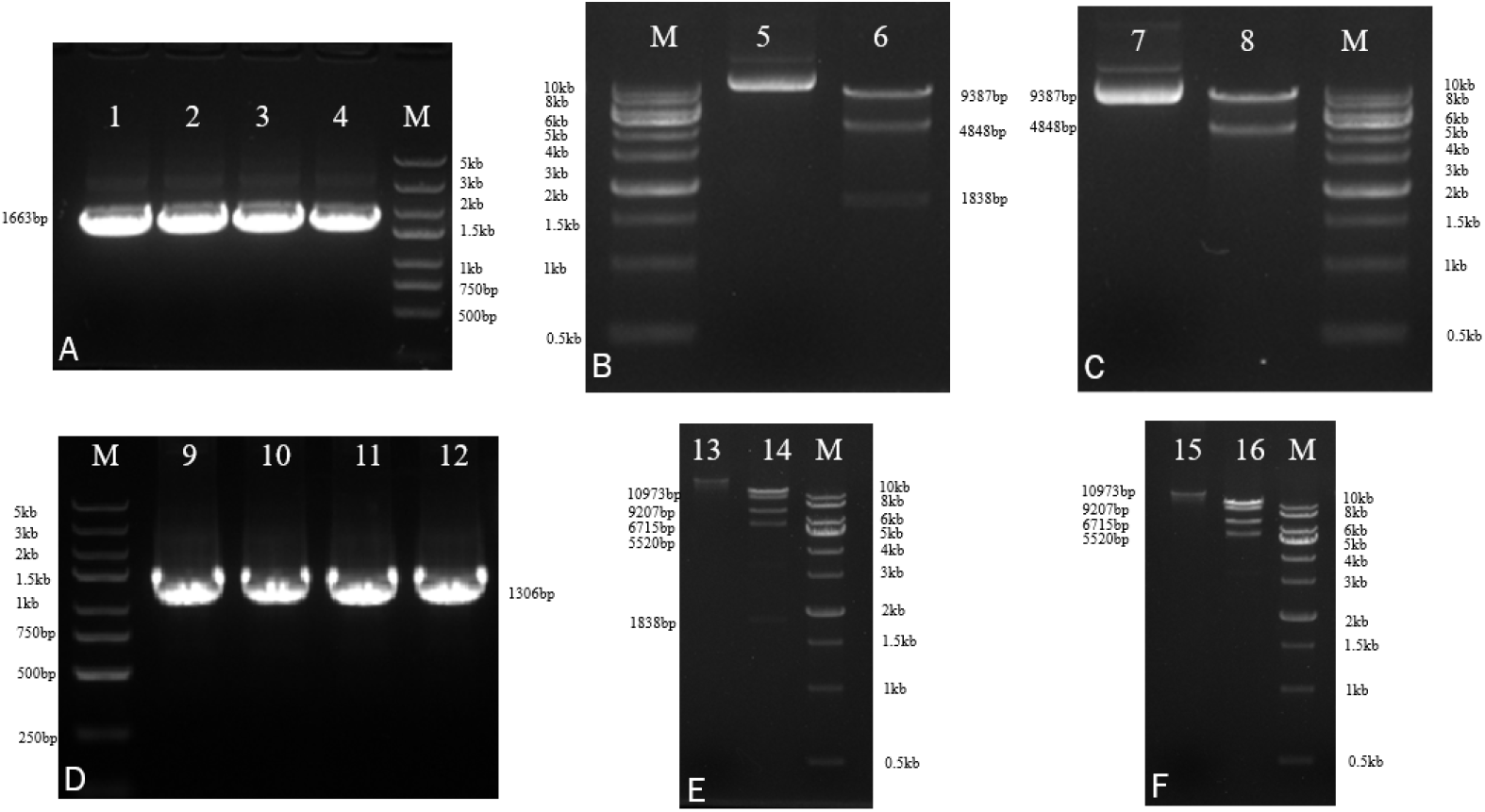
Plasmid yeast universal element knockout verification (A) Colony PCR results of 14 kb plasmid yeast universal element knockout, Lanes 1-4 are the four selected clones, M is 5000bp DNA Marker. (B, C) Enzymatic digestion of 14 kb plasmid, in which lane 5 is the unknocked plasmid, lane 6 is the unknock plasmid digestion results (EcoRI+BamHI), M is the 1 kb DNA Marker, lane 7 is the knocked plasmid, and lane 8 is knockout plasmid digestion results (EcoRI+BamHI), M is a 1kb DNA Marker. (D) Colony PCR results of 32 kb plasmid yeast universal element knockout, Lanes 9-12 are the four selected clones, and M was 5000bp DNA Marker. The enzyme digestion of (E, F) 32kb plasmid verified that lane 13 was the unknocked plasmid, lane 14 is unknock plasmid digestion results(BamhI+XbaI+SpeI), lane 15 is the knocked plasmid, lane 16 is the knockout plasmid digestion results (BamhI+XbaI+SpeI), and M was the 1kb DNA Marker.

**Fig 4.**
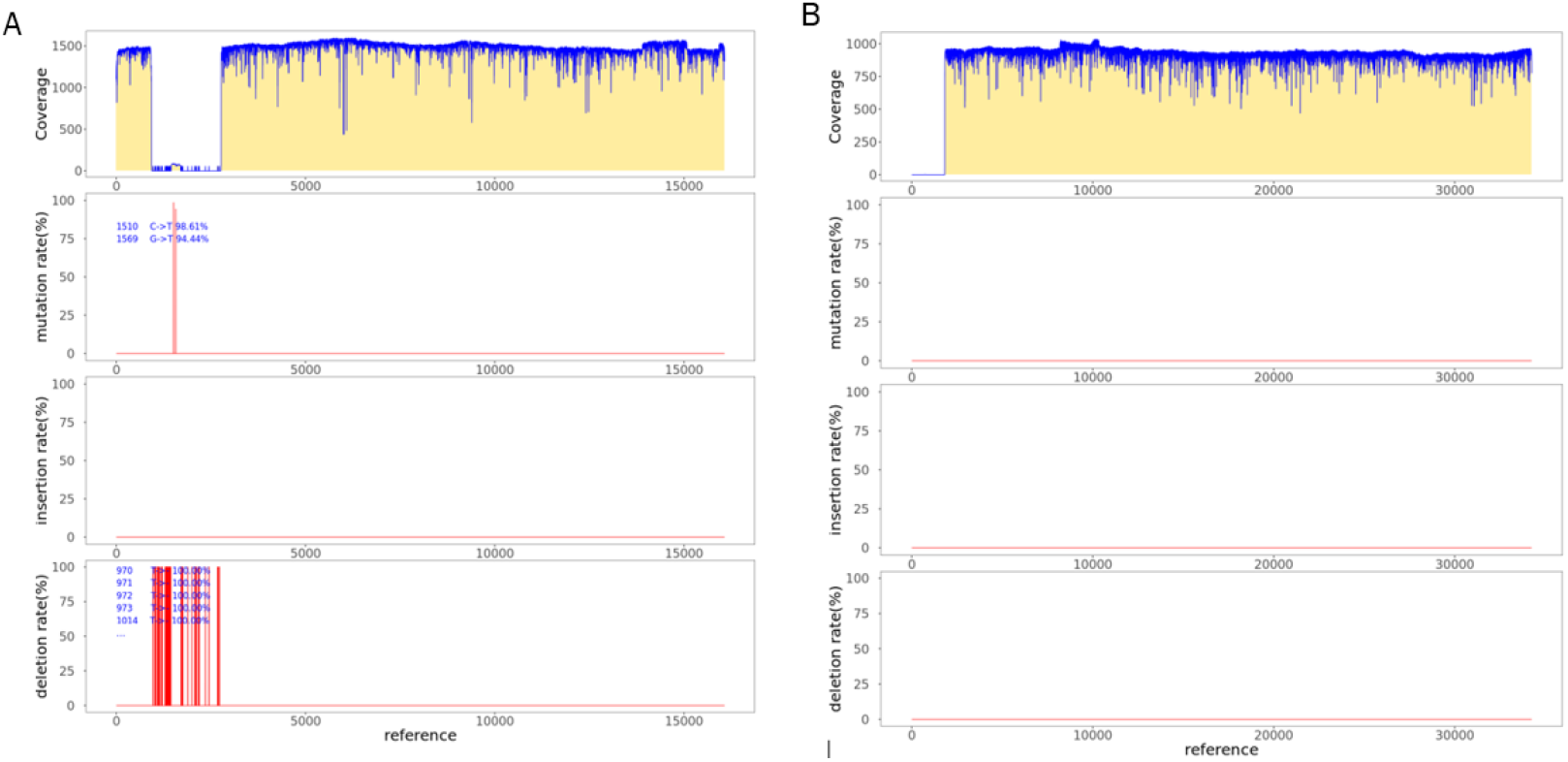
ONT sequencing results of plasmid (A) Analysis of ONT sequencing results of 14kb plasmid (B) analysis of ONT sequencing results of 32kb plasmid

### pCas-SmR plasmid release

In order to obtain a pure target plasmid, we need to remove the tool plasmid pCas-SmR. SacB gene was added when constructing pCas-SmR plasmid. When sucrose was present in the medium, Escherichia coli containing SacB gene plasmid would convert sucrose into fructan, and Escherichia coli itself could not process fructan. This can lead to a large accumulation of fructans and cause the death of the bacteria. According to this principle, the bacteria solution containing the target plasmid and pCas-SmR plasmid will be marked with sucrose plate. Under the survival pressure, E. coli will discard pCas-SmR plasmid, and then a single colony will be coated with spectacular resistant plate to prove the removal of pCas-SmR plasmid. The bacterial solution with two plasmids of 14kb and 32kb constructed in this study could not grow on the plate of spectacomycin after being cultured on the scribe sucrose plate, but could grow normally on the plate of target plasmid resistance, which proved that the pCas-SmR plasmid was successfully removed.

## Materials and method

### STEP 1:Yeast in vivo recombination splicing middle construction

#### Materials

- Saccharomyces cerevisiae BY4741(WEIDI YC1060)
- Zymolyase(SIGMA L4025)
- ChargeSwitch Yeast Plasmid Kit(ThermoFisher CS10203)
- SD-Ura(Solarbio S0620)
- Carrier DNA(WEIDI YC5002)
- PEG/LiAc Solution(WEIDI YC5001)
- DTT(SIGMA DTT-RO)
- glycerin(SIGMA 1295731)

## Methods

1. Preparation of DNA fragments: The full length of plasmid DNA was divided into several fragments, each fragment was equally divided as far as possible, and synthesized by conventional gene synthesis methods. A 50bp homologous repeat region was designed between the fragment and the yeast universal element.
2. Transformation: Each fragment was added to Saccharomyces cerevisiae cell BY4741 at the molar ratio of 1:1, 10μL of Carrier DNA denatured at 96°C was added, 500μL of PEG/LiAc was added, and then the fragment was transferred to 42°C water bath for 30min and then transferred to 42°C water bath for 15min. After centrifuging and discarding the supernatant, the cells were precipitated and re-suspended with 500μL sterile water, then re-suspended with 100μL sterile water, coated on URA defective plates, and incubated at 30°C for 48-96h.
3. Colony PCR verification: Single colonies were selected for further scribing culture, and primers were designed to verify the correctness of the connected region.
4. Yeast Plasmid extraction: The ChargeSwitch Yeast Plasmid Kit was used to extract the yeast plasmid. Take 2-4mL of the yeast solution cultured overnight, centrifuge it at 12000rpm for 2min, discard the supernatant, add 300μL R4 buffer, 10μL Zymolyase (10mg/mL, dissolved in 1xPBS), 20μL 1M DTT, and incubate at room temperature for 1h. Add 300μL L9 buffer to the reaction solution, gently mix it upside down, stand at room temperature for 2min until the solution is clear, add 300μL pre-cooled N5 buffer, gently mix it upside down until flocculent precipitation appears, and centrifuge at 12000rpm for 10min. Mix the supporting magnetic beads well, mix 40μL with the supernatant obtained by centrifugation in the above steps, stand for 1min, and adsorb them on the magnetic rack for 1-2min. Discard the supernatant, wash them once with 1mL W11 and W12, re-suspend the magnetic beads with 50μL E5buffer, incubate them at room temperature for 2min, and adsorb them on the magnetic rack for 1-2min. After the liquid was clarified and clear, 40μL supernatant was absorbed to become yeast plasmid.
5. Preparation of electroreceptor cells: DH10B strain was strewed on LB plate and cultured overnight at 37°C. One monoclone was selected and inoculated into a 4 ml LB liquid medium at 220rpm and 37°C overnight. The 4ml bacterial solution cultured overnight was transferred to 400ml LB medium (v:v=1:100) at 220rpm and cultured at 37°C until OD600=0.6, then the bacterial solution was poured into two 400mL centrifugal bottles and placed on ice for 30-60min. During this period, the bacterial solution was shaken every 20min to quickly and fully cool. Centrifuge at 4000rpm at 4°C for 10min and discard the supernatant. Add about 50ml cold sterile water to each bottle, slowly shake well in ice water, mix 2 bottles into 1 bottle, add cold sterile water to 300ml, shake well, centrifuge at 3500rpm at 4°C for 10min, discard the supernatant. Add about 50ml cold 10% glycerin to each bottle, slowly shake well in ice water, mix 2 bottles into 1 bottle, add cold 10% glycerin to 300ml, shake well, centrifuge at 4000rpm at 4°C for 10min, discard the supernatant. Repeat with 10% glycerin. The cells were re-suspended with 10% glycerol to a final volume of 2-3ml. Finally, the cells were divided into 50μL equal parts in a centrifuge tube and stored at -80°C.
6. Yeast plasmid transformation: Take out the receptive state in the refrigerator at -80°C and thaw it on ice. 5μL yeast plasmid was added into the receptive cells and incubated on ice for 10 min. After the shock was given at 1.8KV, 6ms, 500μL preheated LB was added and cultured at 37°C, resuscitated at 220rpm for 1h, carrier resistant plate was coated and cultured at 37°C overnight.
7. Enrichment of intermediate constructed plasmid: Single colonies on the plate were selected into 4mL LB medium and cultured overnight at 220rpm at 37°C. Plasmid extraction was performed using a commercial plasmid extraction kit.

## STEP2:Removal of yeast universal element

### Materials

- DH10B Chemically Competent Cell WEIDI DL1070)
- L-Arabinose(Solarbio L8060)

### Methods

1. pCas-SmR transformation: DH10B chemoreceptor cells were removed from the refrigerator at -80°C and thawed on ice. 500ng pCas-SmR plasmid was added into receptive cells, mixed gently, and placed in ice bath for 30min. Incubate the mixture in a water bath at 42°C for 60s and immediately place it in an ice bath for 10min. 500μL LB medium was added to the mixture and resuscitated at 220rpm at 37°C for 1 hour. The resuscitated bacterial solution was centrifuged at 5000 RPM for 5min, then coated on a spectacularly resistant LB plate and cultured overnight at 37°C.
2. Preparation of pCas-SmR-DH10B induced electroreceptor state: A monoclonal colony was selected from pCas-SmR-DH10B plate and inoculated into spectacular-resistant LB single tube and cultured overnight at 37°C. 1.5ml bacterial solution cultured overnight was transferred to 150ml LB medium (v:v=1:100), then arabinose with a final concentration of 10mM was added at 220rpm, and when cultured at 37°C to OD600=0.6, 100mL bacterial solution was put into two 50mL centrifugal tubes, and ice bath was performed for 30min. Shake up and down every 20 minutes to cool the bacterial solution quickly and fully. Centrifuge at 4000rpm at 4°C for 10min and discard the supernatant. Add about 10ml sterile water to each tube, mix the bacterial solution well, combine the two tubes into one tube, and add sterile water to a constant volume of 50mL. Centrifuge at 4000rpm at 4°C for 10min and discard the supernatant. Add about 10ml 10% glycerin, mix the bacterial solution well and add 10% glycerin to 50ml. Centrifuge at 4000rpm at 4°C for 10min and discard the supernatant. Wash again with 10% glycerin water. Add 1mL of 10% glycerin water to the suspenseful bacterial solution, pack it into a centrifuge tube at 50μL per tube, and store it at -80°C.
3. Donor DNA preparation: Primers are designed to amplify the 450 bp DNA fragments adjacent to the yeast universal element, and the two PCR products are combined into a 900bp length Donor DNA by fusion PCR, which serves as a template for homologous recombination.
4. Electrical transformation: Take pCas-SmR-DH10B receptive cells out of the refrigerator at -80°C, melt them on ice, add 500ng intermediate plasmid containing yeast universal element and 200ng Donor DNA, gently knock and mix them, and incubate them on ice for 10 min. After electric shock at 1.8KV, 6ms, 500μL preheated LB was added to culture at 37°C, resuscitated at 220rpm for 1h, coated with carrier resistance and spectinomycin resistance plate, and cultured at 37°C overnight.
5. Liquid PCR to verify the knockout of yeast universal elements: Single colony shake bacteria on the plate were selected, and primers were designed to verify the knockout of yeast universal elements through liquid PCR.

## STEP3:Deletion of pCas-SmR plasmid

### Materials

- Glucose (WEIDI C1010-01)
- Sucrose(Solarbio L8271)

### Methods

1. The bacteria solution containing the final plasmid was coated with LB plate with sucrose concentration of 10g/L and glucose concentration of 5g/L, and cultured at 37°C overnight.
2. Single colony was selected and shaken overnight at 37°C, 220rpm.
3. The bacterial solution is coated with carrier resistant and spectinomycin resistant plates. Clones that grow on carrier resistant plates but do not grow on spectinomycin plates are positive clones.
4. 100μL of successfully knocked out bacterial solution was transferred to 4 ml carrier resistant LB medium for overnight culture at 37°C. The plasmid was extracted by commercial kit and verified by enzyme digestion and third-generation sequencing.

## Discussion

In recent years, DNA synthesis technology has been continuously developed, the length of synthetic DNA has been continuously improved, and the synthetic assembly technology has been continuously innovated. The DNA assembly technology of small fragments has been relatively stable, while the DNA assembly of large fragments is not suitable for in vitro assembly technology due to its large molecular weight and easy to break, and it usually needs to adopt the biological splicing scheme. In this study, saccharomyces cerevisiae cells were chosen as the assembled host cells because of their better recombination ability and stability compared with E. coli and Bacillus subtilis, etc. The results of this paper also showed that the success rate of using yeast cells for splicing reached 100%. As the third generation gene editing technology, CRISPR-Cas9 gene editing technology has the advantages of high editing efficiency, simplicity and speed, and is widely used in gene therapy, biological modification and other directions. In this study, CRISPR-Cas9 system is used to carry out gene knockout of large plasmids. The results showed that the success rate of CRISPR-Cas9 system on yeast universal element was 100%, showing good success rate and stability.

In order to get rid of the requirement of in vivo yeast splicing technology for plasmid yeast elements, we proposed a large fragment DNA splicing scheme combining in vivo yeast splicing technology and CRISPR-Cas9 technology. Compared with the traditional scheme, this scheme does not need to rely on yeast plasmid, and can realize the splicing of any carrier in yeast by using the yeast universal element selected in this study, and can splicing 30kb DNA fragments efficiently. At the same time, the yeast universal element in the spliced vector can be effectively knocked out, and the original vector sequence can be maintained without modifying the vector sequence, which has a broad application prospect in the field of constructing long DNA fragments. At the same time, there are some disadvantages in the assembly of large fragments of DNA using in vivo splicing technology. For example, the recombinant arm cannot be homologous with the yeast genome, otherwise it will be recombined with the yeast genome. Some cytotoxic genes also cannot be assembled in this way; Sequences with complex structure, such as Poly structure, high GC, dense repetition, etc., will affect the success rate of assembly. Therefore, yeast homologous recombination technology still needs to be further optimized in the future.

## Supporting information

SI Tabl: Full-length sequence of each plasmid

S2 Tabl: primer information

S3 Tabl: Sequencing results of 14KB plasmid ONT

S4 Tabl:32KB plasmid ONT sequencing results

## Acknowledgments

Thanks to Azenta’s’ foundation for supporting this study.

## Supplement

SI Tabl: Full-length sequence of each plasmid

S2 Tabl: primer information

S3 Tabl: Sequencing results of 14KB plasmid ONT

S4 Tabl:32KB plasmid ONT sequencing results

## References

1. Lartigue C, Vashee S, Algire MA, et al. Creating bacterial strains from genomes that have been cloned and engineered in yeast. Science. 2009;325(5948):1693–1696. doi:10.1126/science.1173759

2. Ma Y, Zhang L, Huang X. Genome modification by CRISPR/Cas9. FEBS J. 2014;281(23):5186–5193. doi:10.1111/febs.13110

3. Hryhorowicz M, Lipiński D, Zeyland J, Słomski R. CRISPR/Cas9 Immune System as a Tool for Genome Engineering. Arch Immunol Ther Exp (Warsz). 2017;65(3):233–240. doi:10.1007/s00005-016-0427-5

4. Annaluru N, Muller H, Mitchell LA, et al. Total synthesis of a functional designer eukaryotic chromosome [published correction appears in Science. 2014 May 23;344(6186):816]. Science. 2014;344(6179):55–58. doi:10.1126/science.1249252

5. Cello J, Paul AV, Wimmer E. Chemical synthesis of poliovirus cDNA: generation of infectious virus in the absence of natural template. Science. 2002;297(5583):1016–1018. doi:10.1126/science.1072266

6. Kosuri S, Church GM. Large-scale de novo DNA synthesis: technologies and applications. Nat Methods. 2014;11(5):499–507. doi:10.1038/nmeth.2918

7. Engler C, Marillonnet S. Golden Gate cloning. Methods Mol Biol. 2014;1116:119–131. doi:10.1007/978-1-62703-764-8_9

8. Li L, Lu Y, Jiang W. Sheng Wu Gong Cheng Xue Bao. 2013;29(8):1113–1122.

9. Gibson DG, Smith HO, Hutchison CA 3rd, Venter JC, Merryman C. Chemical synthesis of the mouse mitochondrial genome. Nat Methods. 2010;7(11):901–903. doi:10.1038/nmeth.1515

10. Gibson DG, Glass JI, Lartigue C, et al. Creation of a bacterial cell controlled by a chemically synthesized genome. Science. 2010;329(5987):52–56. doi:10.1126/science.1190719

11. Gunge N. Yeast DNA plasmids. Annu Rev Microbiol. 1983;37:253–276. doi:10.1146/annurev.mi.37.100183.001345

12. Jazwinski SM. Replication of the 2-micrometer DNA plasmid of yeast. Acta Biochim Pol. 1982;29(1-2):159–173.

13. Gupta D, Bhattacharjee O, Mandal D, et al. CRISPR-Cas9 system: A new-fangled dawn in gene editing. Life Sci. 2019;232:116636. doi:10.1016/j.lfs.2019.116636

14. Gostimskaya I. CRISPR-Cas9: A History of Its Discovery and Ethical Considerations of Its Use in Genome Editing. Biochemistry (Mosc). 2022;87(8):777–788. doi:10.1134/S0006297922080090

15. Bao A, Burritt DJ, Chen H, Zhou X, Cao D, Tran LP. The CRISPR/Cas9 system and its applications in crop genome editing. Crit Rev Biotechnol. 2019;39(3):321–336. doi:10.1080/07388551.2018.1554621

16. Flagg MP, Kao A, Hampton RY. Integrating after CEN Excision (ICE) Plasmids: Combining the ease of yeast recombination cloning with the stability of genomic integration. Yeast. 2019;36(10):593–605. doi:10.1002/yea.3400

17. Fitzgerald-Hayes M. Yeast centromeres. Yeast. 1987;3(3):187–200. doi:10.1002/yea.320030306

18. Boeke JD, Trueheart J, Natsoulis G, Fink GR. 5-Fluoroorotic acid as a selective agent in yeast molecular genetics. Methods Enzymol. 1987;154:164–175. doi:10.1016/0076-6879(87)54076-9

19. Chen W, Zhang Y, Zhang Y, et al. CRISPR/Cas9-based Genome Editing in Pseudomonas aeruginosa and Cytidine Deaminase-Mediated Base Editing in Pseudomonas Species. iScience. 2018;6:222–231. doi:10.1016/j.isci.2018.07.024

20. Li Q, Sun B, Chen J, Zhang Y, Jiang Y, Yang S. A modified pCas/pTargetF system for CRISPR-Cas9-assisted genome editing in Escherichia coli. Acta Biochim Biophys Sin (Shanghai). 2021;53(5):620–627. doi:10.1093/abbs/gmab036

21. Gietz RD, Schiestl RH. High-efficiency yeast transformation using the LiAc/SS carrier DNA/PEG method. Nat Protoc. 2007;2(1):31–34. doi:10.1038/nprot.2007.13

